# Targeted memory reactivation is not more effective during slow wave sleep than sleep stage 2

**DOI:** 10.1101/2023.10.19.563121

**Authors:** Julia Carbone, Carlos Bibian, Jan Born, Cecilia Forcato, Susanne Diekelmann

## Abstract

Sleep facilitates memory consolidation, which is assumed to rely on the reactivation of newly encoded memories orchestrated by the temporal interplay of slow oscillations (SO), fast spindles and ripples. SO as well as the number of spindles coupled to SO are more frequent during slow wave sleep (SWS) compared to lighter sleep stage 2 (S2). But, it is unclear whether memory reactivation is more effective during SWS than during S2. To test this question, we applied Targeted Memory Reactivation (TMR) by presenting learning-associated sound cues during SWS vs. S2 in a counterbalanced within-subject design. Contrary to our hypothesis, memory performance was not significantly better when cues were presented during SWS. Event-related potential (ERP) amplitudes were significantly higher for cues presented during SWS than S2, and the density of SO and SO-spindle complexes was generally higher during SWS than during S2. Whereas SO density increased during and after the TMR period, SO-spindle complexes decreased. None of the parameters were associated with memory performance. These findings suggest that the efficacy of TMR does not depend on whether it is administered during SWS or S2, despite differential processing of memory cues in these sleep stages.

## Introduction

Sleep is widely assumed to support memory consolidation^1–4^. The beneficial function of sleep for memory has been proposed to rely on the reactivation of newly encoded memory traces during subsequent sleep. During a night of sleep, the brain undergoes different sleep stages that alternate in a cyclic manner. The different sleep stages may play distinct roles in memory reactivation and consolidation^5^. Human sleep is composed of rapid eye movement (REM) sleep and non-REM (NREM) sleep, which includes light sleep (stages S1 and S2) and deep sleep, i.e., slow wave sleep (SWS, stages S3 and S4). An increasing number of studies implicates mainly NREM sleep in the reactivation and consolidation of declarative memories, i.e., memory for facts and events^6^. NREM sleep is characterized by specific electrophysiological brain oscillations, particularly neocortical slow oscillations (SO, 0.5-1 Hz), thalamocortical spindles (9-15 Hz), and hippocampal ripples (80–200 Hz)^7–9^. The precise temporal coordination between these oscillations is assumed to constitute a crucial mechanism for memory consolidation^10–12^. SO provide a temporal frame for the occurrence of spindles mainly in the excitatory up-state of the SO, with ripples in turn being preferentially nested in the spindle troughs^9^. This temporal coupling orchestrates the reactivation of hippocampal memory representations to promote their redistribution and integration into neocortical networks for long-term storage^1,13–15^.

Although there is convincing evidence of the importance of these mechanisms for the reactivation and consolidation of memory, it is still a matter of debate whether the different sleep stages of NREM sleep, particularly S2 and SWS, play distinct roles in these processes^1,16^. SO are evident during both S2 and SWS, but to a much higher degree in SWS^17^. In fact, the amount of slow wave activity (SWA, 0.5-4 Hz) constitutes the main sleep feature to distinguish between S2 and SWS, with more than 20% of slow waves per sleep epoch indicating SWS according to standard criteria by Rechtschaffen & Kales^18^. A number of studies found the amount and intensity of SWS as well as of SWA and SO to be associated with better memory consolidation^19–22^. Sleep spindles are present during both S2 and SWS to a similar degree, and stage 2 spindle activity was also found to be related to memory consolidation^23–26^. Since S2 covers a higher percentage of the sleep night, some researchers pointed out the relevance of lighter sleep stage S2 for memory consolidation^16,24^. Others have argued that it is mainly the temporal coupling of spindles to SO that is essential for the reactivation of memories during sleep, and since there are more SO and more frequent SO-spindle coupling during SWS^27^, SWS could be assumed to be more important for the processing of memories than S2.

In the present study, we directly compared the effectiveness of memory reactivation and consolidation during SWS and S2. To experimentally manipulate memory reactivation during specific sleep stages, we applied a technique known as targeted memory reactivation (TMR)^28,29^. In TMR, specific stimuli like odors or sounds become associated with learning contents during the encoding phase, and the same stimuli are then presented again as learning-associated cues during subsequent sleep. TMR is assumed to trigger and/or facilitate endogenous processes of memory reactivation and has consistently been shown to stabilize and enhance declarative memory^28–31^. Previous studies using TMR showed that the electrophysiological signatures of memory reactivation are characterized by a temporary increase in theta activity (4-8 Hz) as well as of spindle activity after re-exposure to the cues during NREM sleep^32,33^. Many previous studies on TMR during sleep presented reminder cues exclusively during SWS^31,34–38^. Others have applied TMR during the entire NREM sleep period^32,39–42^ or during the entire night irrespective of sleep stage^43–46^. According to a recent meta-analysis, TMR was effective during both S2 and SWS, although there were only very few studies for TMR applied only during S2 compared to a large number of studies on TMR during SWS or S2+SWS^28^. Importantly, this meta-analysis did not differentiate between cueing that took place exclusively during SWS and cueing during both S2+SWS.

In the present study, we directly compared TMR applied during S2 with TMR during SWS for declarative memory consolidation in the same participants. Based on the assumption that sleep-dependent memory reactivation strongly relies on the temporal coupling of spindles and ripples to SO, which is more pronounced during SWS, we hypothesized that TMR would be more effective during SWS than S2.

## Results

Participants learned 30 sound-word associations in the evening, with the sounds and words being semantically related (e.g., the sound of coins dropping plus the German word for ‘casino’)^37,48^ (Figure 1a). During subsequent polysomnographically recorded sleep, auditory cues (sounds plus the first syllable of the words) were presented via earphones, with half of the cues being presented during SWS, and the other half during S2 (in counterbalanced order) during the first sleep cycle (Figure 1b). Cued recall was tested in the next morning after learning of an interference task. Memory performance was assessed as ‘memory change’ from training to testing to control for individual differences at learning (i.e., correct responses at testing minus correct responses at training). 20 subjects were included in the final analyses (see methods section for exclusion criteria).

**Figure 1.**
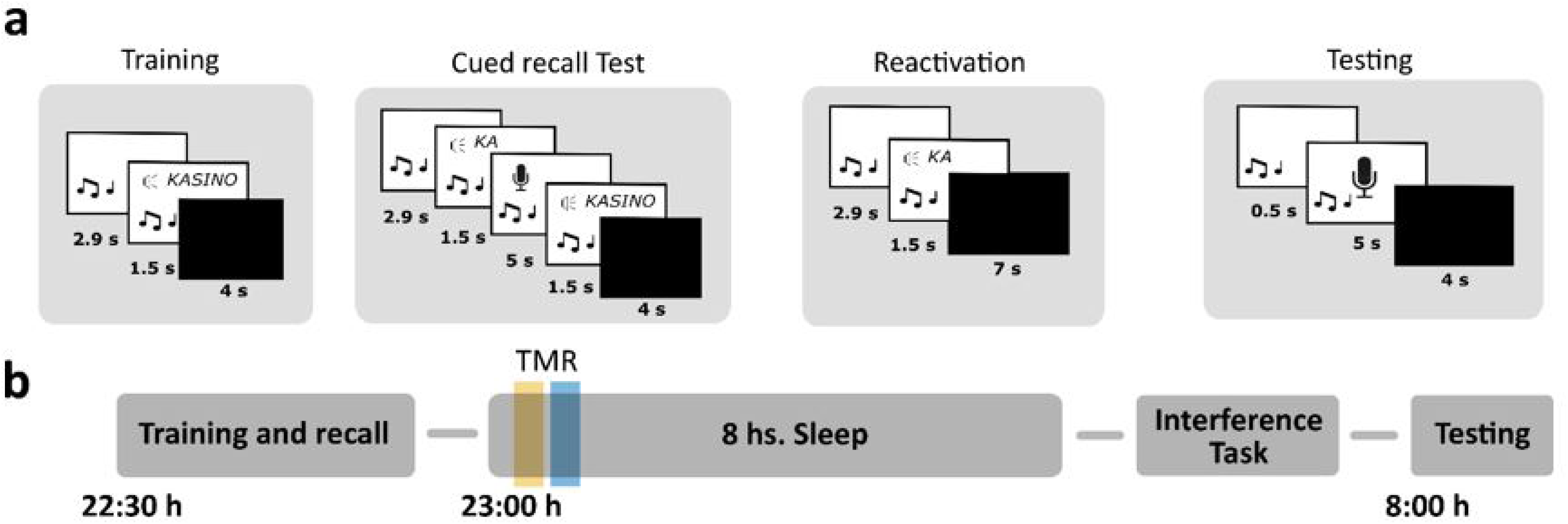
Memory task and experimental design. a. During training, 30 sound-word associations were presented (German words). For each association, the sound was presented first for 2.9 s and continued accompanied by the word written on the screen and spoken aloud for 1.5 s. After a 4 s break, the next association appeared. In the cued recall test, for each association, the sound was presented for 2.9 s and continued accompanied by the first syllable of the associated word for 1.5 s. Afterwards, a microphone appeared on the screen and participants had 5 s to say the word aloud (sound continued during the entire period). Independently of their answer, correct feedback was given with the word spoken and written on the screen for 1.5 s. During Targeted memory reactivation (TMR, Reactivation), each sound was first presented alone for an average of 2.9 s, then the syllable was played once with the sound continuing in the background for another 1.5 s. After a 7 s break, the next cue was presented (until each cue was presented once). In the interference task, subjects performed the same task as for the training session with the same sounds associated to new words. During testing, each sound was presented for 0.5 s and then subjects were asked to say the word aloud. **b.** Training and the cued recall test took place in the evening (22:30 h) and subjects went to bed at ∼23:00 h. TMR took place during the first sleep cycle, with half of the cues being presented in SWS and the other half in S2 (orange and blue lines) in counterbalanced order. In the next morning, subjects learned an interference task and took part in the testing session.

### Memory performance

To assess the differences in memory performance between cues presented in S2 and SWS, we performed a repeated-measures ANOVA with the within-subjects factor “sleep stage” (S2 vs. SWS) and the between-subjects factor “order” (first TMR in S2 vs. first TMR in SWS). Results revealed that memory performance was not more effective during SWS than S2 (Figure 2a). Performance was even slightly but non-significantly better for S2 cueing (main effect sleep stage: F_1,18_ = 4.05, P = 0.059). The order of conditions did not affect memory performance (main effect order: F_1,18_ = 0, P = 1; interaction effect: F_1,18_ = 2.45, P = 0.135). When comparing memory change directly between SWS and S2 (not taking order of conditions into account), similar results were obtained (P = 0.067, Table 1). The study yielded a statistical power of 0.83 for a one-tailed comparison (given our hypothesis of S2 < SWS), with an effect size of 0.6 and a sample size of 20, indicating a relatively high likelihood of detecting the specified effect in the analysis. Given the absence of significant differences between conditions, we formally tested whether our data support the null hypothesis of SWS not providing better cueing effects than S2. A Bayesian paired t-Test of our original hypothesis (i.e., S2 < SWS) using a standard prior (Cauchy distribution with scale factor 0.707) yielded strong evidence for the null model assuming equivalent performance between conditions (BF_01_ = 11.06; see Supplementary Figure S1). This was confirmed by robustness checks across different prior widths, as well as a sequential analysis across participants.

**Figure 2.**
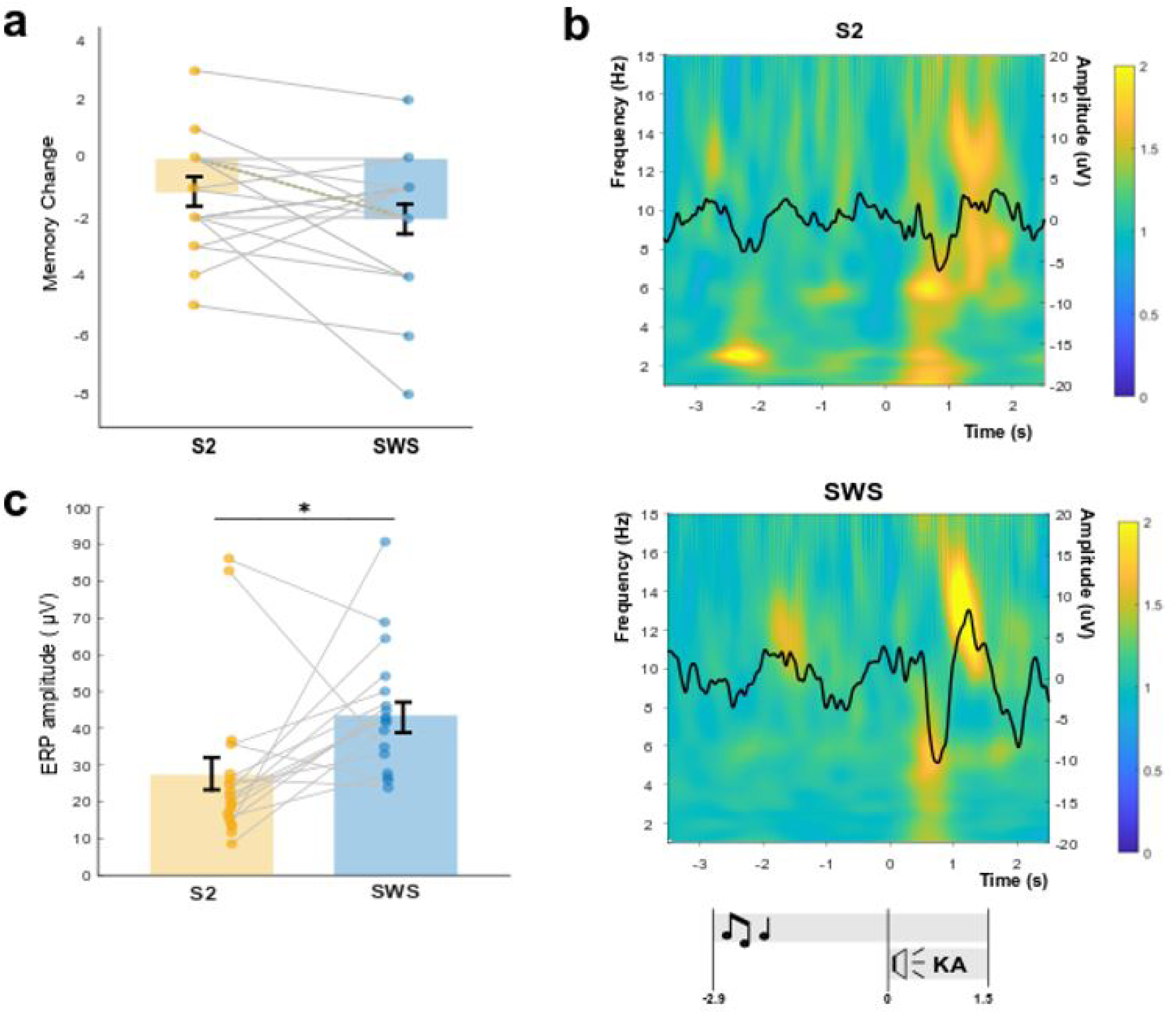
Memory performance and changes in oscillatory activity upon cueing during SWS and S2. a. Participants’ performance was not significantly different for cues presented in S2 or SWS. Memory change: number of correct words at testing minus number of correct words at training. Means ± SEM are shown. **b.** Time–frequency representations (TFR) for cues presented during S2 and SWS, each with their corresponding event-related potential (ERP). Color maps show power changes relative to a 1 s baseline right before sound onset (i.e. −4 to −3 s). TFR were aligned at time-point zero to the syllable cue onset. Lower panel indicates timeline for the sound (symbolized by musical notes) and the syllable cue (symbolized by speaker and ‘KA’ syllable). **c.** Cues presented in SWS elicited larger ERP amplitudes than those in S2. Average ERP amplitudes for single subjects as well as the means ± SEM are shown. * P < 0.05.

**Table 1.** Memory measures.

|  | *Overall* | *Cues presented in S2* | *Cues presented in SWS* | *P values S2 vs. SWS* |
| --- | --- | --- | --- | --- |
| Training | $22.9 \pm 1.0$ | $11.0 \pm 0.6$ | $11.9 \pm 0.4$ | 0.025 |
| Testing | $19.7 \pm 1.5$ | $9.9 \pm 0.8$ | $9.8 \pm 0.8$ | 0.9 |
| Memory change | $-3.2 \pm 0.8$ | $-1.15 \pm 0.4$ | $-2.05 \pm 0.5$ | 0.067 |
| Interference task - Training | $24.5 \pm 0.8$ | $12.4 \pm 0.4$ | $12.1 \pm 0.5$ | 0.6 |
| Interference task - Testing | $21.5 \pm 1.2$ | $10.6 \pm 0.7$ | $10.9 \pm 0.7$ | 0.6 |
| Interference task - Memory change | $-3.0 \pm 0.7$ | $-1.8 \pm 0.4$ | $-1.3 \pm 0.5$ | 0.4 |

Number of correct responses at training and testing for the original task and the interference task (means ± SEM). Memory change refers to correct responses at testing minus correct responses at training. P values are indicated for paired sample t-tests.

Initial learning during the Training session differed unexpectedly for S2 and SWS cues (P = 0.025, Table 1), which was controlled for by calculating the memory change from Training to Testing. Performance for the Interference Task (i.e., same sounds associated to new words) was comparable between S2 and SWS cues for the Training session (P = 0.6), Testing session (P = 0.6) and for memory change (P = 0.4, Table 1).

To control for possible awareness of cue presentation during sleep, in the end of the experiment, subjects were presented with all reminder cues again and were asked to report whether they think that they have heard the cues during the night (‘Heard/not-heard task’). On average, they correctly recognized 6.2 ± 1.5 cues, with no difference between the SWS and S2 cueing conditions (P = 0.26).

### Sleep analyses

Sleep scoring was performed according to standard criteria by Rechtschaffen & Kales^18^. All subjects included in the final analysis showed normal sleep patterns, with a total sleep time of 490.0 ± 3.1 min, and average time spent in the sleep stages (in min, mean ± SEM) of S1 (15.4 ± 2.1), S2 (290.4 ± 7.4), S3 (65.1 ± 5), S4 (13.8 ± 2.4), REM sleep (87.8 ± 6.3) and wake (19.9 ± 8.8).

To test for differences in sleep oscillatory activity when cues were presented during S2 and SWS, we performed time-frequency analyses upon cue presentation separately for S2 and SWS (Figure 2b). First, we looked at the event-related potentials (ERP) upon the onset of the syllable cue, considering that the syllable (and not the sound) was found to be the relevant cue triggering memory reactivation in previous studies^37,48^. In both sleep stages, there was an elicited response upon syllable cue onset, however, the ERP amplitude was significantly higher when cues were presented in SWS compared to S2 (P = 0.012) (Figure 2c).

Next, we compared the density of slow oscillations (SOs), fast spindles, and SO-spindle complexes (i.e., SO and fast spindles occurring together) during three periods of interest: the entire period of cue presentation (“React”, 3 min for 15 cues), a pre-cueing period immediately before the cueing period (“Pre”, 3 min), and a post-cueing period immediately after the end of the cueing period (“Post”, 3 min). Repeated-measures ANOVAs (“sleep stage” x “Pre/React/Post”) revealed significant interaction effects for all three measures (all P < 0.002). SO density in SWS significantly increased from Pre to React (P < 0.001) and from Pre to Post (P < 0.001, Figure 3). Moreover, when comparing equivalent periods in S2 and SWS, SO density was significantly higher in SWS for all time windows (all P < 0.001). Spindle density was significantly higher during the Pre period in SWS than in S2 (P = 0.007), while a spindle density increase was observed from Pre to Post in S2 (P = 0.001). The density of SO-fast spindle complexes followed a similar pattern as SO density, with a higher density of SO-spindle complexes in SWS compared to S2 (all P < 0.001). However, during SWS, the density of SO-spindle complexes decreased from Pre to React (P < 0.001), and from Pre to Post (P < 0.001). There were no significant correlations between any of these measures and memory change. Bayesian correlations yielded moderate evidence for the null model (3 < BF_01_ < 4).

**Figure 3.**
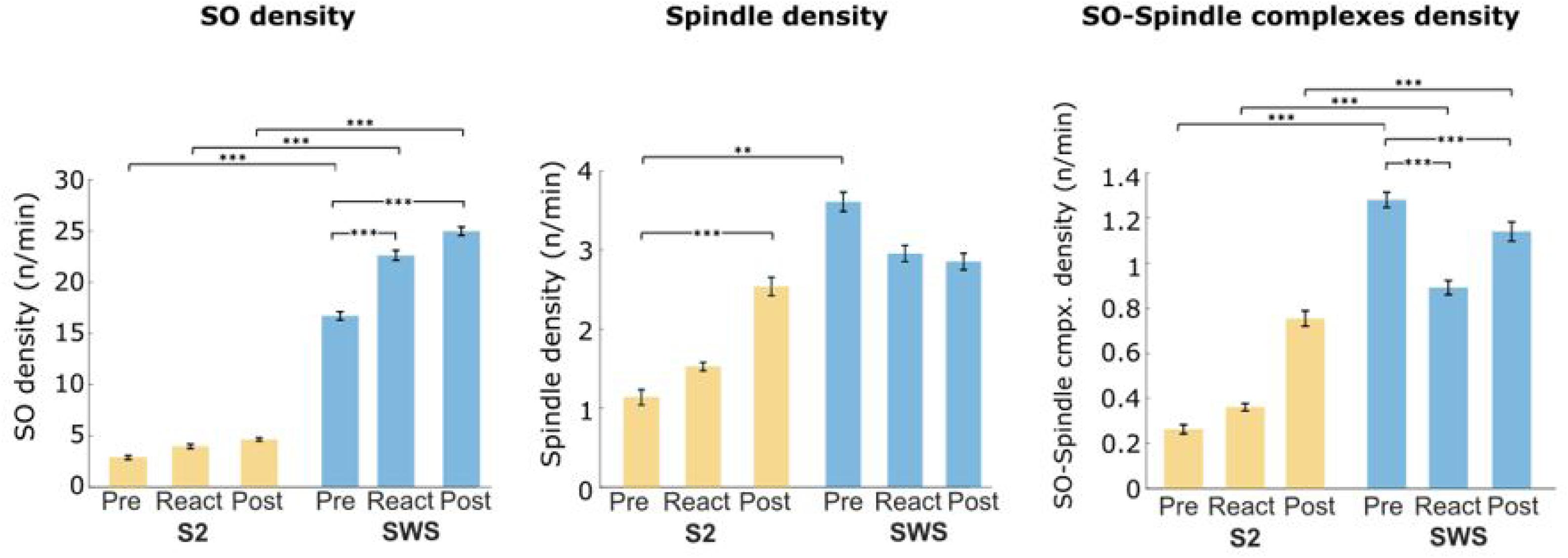
**Density of slow oscillations (SO), fast spindles and SO-Spindle complexes**. Each bar indicates density calculated within a period of interest: “Pre”, a 3-min period immediately before the cueing period; “React”, during the 3-min cueing period; and “Post”, a 3-min period immediately after the cueing period. Yellow bars correspond to S2 and blue bars to SWS. SO density and SO-spindle complexes density are shown for central electrodes, while fast spindle density is shown for parietal electrodes. Means ± SEM are shown. ** P < 0.01, *** P < 0.001.

## Discussion

In the present study, we compared the effectiveness of targeted memory reactivation (TMR) for the strengthening of declarative memories during the two main NREM sleep stages, S2 and SWS. Our results suggest that, at least for the time intervals analyzed between reactivation and testing in this study, TMR is not more effective during SWS, but both light and deep NREM sleep offer comparably good windows of opportunity for TMR. We explored reactivation-related brain activity for each sleep stage, confirming the expected electrophysiological responses upon cueing during NREM sleep. Furthermore, we observed higher ERP amplitudes for the cue-evoked responses during SWS compared to S2. Reactivation during SWS induced increases in slow oscillation (SO) density and decreases in the density of SO-fast spindle complexes during the reactivation period and thereafter. Reactivation during S2, on the other hand, induced increases in fast spindle density when comparing periods before and after reactivation. Overall, the density of SO and SO-fast spindle complexes was higher during SWS when compared to equivalent periods in S2. Interestingly, the reactivation-induced changes in oscillatory patterns were not correlated with memory performance.

The finding that TMR during SWS is not better than during S2 was in contrast with our hypothesis; we expected higher memory performance when TMR was performed during SWS than during S2. The findings suggest that, overall, subjects showed forgetting of the sound-word associations, and this was consistent for cueing during S2 and SWS. Previous studies argued that declarative memory consolidation is associated with SWS and processes characteristic of SWS, such as SO as well as the coupling between SO and spindles^7,19–21,49^. However, others have argued that lighter sleep stage S2 may be more important for active processes of memory reactivation than SWS^16^. Our results speak in favour of the importance of both NREM sleep stages, at least concerning externally triggered reactivation processes in the time frame examined here. Although any inference with regard to spontaneous reactivation during sleep remains tentative, we conclude that the brain seems to be equally susceptible for the induction of reactivation processes triggered by external memory cues in SWS and S2. This conclusion is in line with recent findings showing no differences in memory performance when TMR was performed during S2 and SWS for a vocabulary task^47^.

In the present study, we even observed descriptively stronger effects of TMR during S2, which should be explored in future research. Although we cannot say whether TMR may be more effective during S2, our results provide convincing evidence against our original hypothesis of stronger TMR effects in SWS. Effect size analyses together with Bayesian statistics indicate a relatively high likelihood of detecting the specified effect in the analyses and provide positive evidence for the null results.

Inspection of EEG activity for cueing during SWS and S2 confirmed the expected electrophysiological responses upon TMR during NREM sleep^32,50,51^. In both sleep stages, time frequency representations showed that there was an evoked response upon the presentation of the cue syllable characterized by a negative peak with power increases in theta/slow spindle frequencies and a positive peak with increased power in the fast spindle band. These results confirm that cues were processed in the brain in both sleep stages. Interestingly, the ERP amplitude was significantly higher during SWS than S2, although it is unclear whether higher ERP amplitudes reflect better responses towards cueing during sleep. There are findings showing that the amplitude of SO, akin to ERPs, is generally higher during SWS relative to S2^17^, and here we did not find an association between ERP amplitude and memory performance. Thus, it remains to be elucidated whether the stronger increase in ERP amplitude during SWS is functionally related to cue-elicited memory processing.

When TMR was applied during SWS, we observed an increase in SO density during the cueing period as well as shortly after the cueing period, when compared to a pre-cueing period of equivalent length. SO are the main feature of SWS, with about four times as many SO during SWS than during S2^17^. Interestingly, this pattern of increase from pre to post cueing was not observed for S2, despite the presence of ERPs. On the other hand, when looking at the density of fast spindles during the same periods, we found a significant increase from pre to post cueing during S2 but not in SWS. These observations are in line with previous studies mainly implying SO for the effects of SWS^19–22^ and sleep spindles for the effects of S2 for declarative memory consolidation^23–26^. Surprisingly, the temporal coupling between SO and spindles, which is assumed to be functionally related to memory consolidation during sleep^13,14,27,52^, decreased from before to after the cueing period in SWS. This finding may explain the lack of stronger cueing effects during SWS compared to S2. However, considering that none of the observed changes in electrophysiological responses were associated with memory performance, the functional relevance of these responses remains to be elucidated.

Our study has some limitations. First, the memory task and the reactivation protocol were mainly designed to test our behavioral hypothesis but were not ideally suited for EEG analyses. We used a low number of reactivation cues, i.e., just one repetition of each of the 30 associations, which limited the power for the ERP and time-frequency analyses. Second, the sound-syllable cues of 3.5 s length were played upon detection of stable S2 or SWS. This means that TMR was not time-locked to endogenously detected SO, neither to a specific phase of the SO. The length of the cues was also longer than an average SO, in some cases leading to more than one evoked response per trial. Another limitation is related to the fact that our SO analyses, i.e., SO density and SO-spindle complexes, resulted in low numbers or even values of zero during S2 in some cases, due to the low number of SO in S2 for the short reactivation periods. Moreover, we observed differences in learning levels for cues to be presented in SWS vs. S2. Although we corrected for this difference by calculating the memory change from training to testing, the findings should be interpreted with caution. Finally, our study did not include a control condition without TMR. Thus, we cannot say whether cueing during S2 and SWS was effective per se, when compared to an appropriate control condition. This should be subject to further investigation.

## Methods

### Participants

A total of 29 participants took part in the study. Sample size was based on power analyses from our previous experiments using the same memory task^37,48^. Nine participants were excluded from the final analysis due to an incorrect audio recording (3 subjects), an incomplete reactivation session (2 subjects), an incomplete performance of the memory task (1 subject), or an extended period of wakefulness between the auditory cues (2 subjects) or during the night (1 subjects), resulting in 20 remaining subjects (M_age_ = 21.6 ± 0.5, 13 female). Participants reported having a regular sleep-wake cycle and not carrying out shift work at least 6 weeks before the experiments. They further did not have any history of neurological, psychiatric or endocrine disorder, did not take any medication at the time of the experiments, and were non-smokers. All participants had an adaptation night and gave informed consent before participation in the experiment. Participants received a financial compensation for their participation in the study. The study was approved by the ethics committee of the Medical Faculty of Tübingen University.

### General procedure and design

The experiment started in the evening at ∼21:30 h with placement of the electrodes for polysomnographic recording. The training session for the sound-word association memory task started at ∼22:30 h (Figure 1b). Directly afterwards, subjects slept 8 h with targeted memory reactivation during the first sleep cycle. In the next morning, subjects learned an interference task and, thereafter, were tested for both tasks.

### Memory task

The task consisted of 30 associations between sounds and semantically related German words (for example, the sound of dropping coins associated with the word KASINO [casino]). Each sound had a duration of between 2,855 and 2,940 ms (on average 2.9 s). All words had three syllables and were pre-recorded by a female voice.

*Training session*: Each trial started with the sound presentation for 2.9 s, followed by the presentation of the associated word written on the screen and spoken aloud once via headphones, meanwhile the sound continued being played in the background for 1.5 s (see Figure 1a). After 4 s break, the next association appeared. After all 30 associations were presented once, subjects performed a cued recall test to get an immediate measure of learning. For each association, subjects listened to the sound and the first syllable of the associated word, and were asked to say the complete word aloud. After 5 s, and independently of their answer, the sound plus the correct word was presented both written on the screen and aloud via headphones to provide correct feedback (1.5 s). Subjects that did not reach 40% correct responses (12 correct answers) during immediate recall were excluded from the analysis.

*Interference task:* The same sounds from the training session were associated with new words for the interference task. The words were also semantically related to the sounds and likewise consisted of three syllables, but with a different first syllable than the words of the original task. The procedure was exactly the same as for the training session.

*Testing session*: For each sound-word association, the sound was played for 0.5 s while a microphonés image appeared on the screen notifying the subjects to say the associated word aloud. After a break of 4 s, the procedure continued with the next sound until all 30 associations were presented.

### Targeted memory reactivation

Participants were asked to wear earphones just before going to sleep and to confirm they could hear white noise (43 dB), which was presented from going to bed until the reactivation session finished. Cueing was performed during the first hour of sleep: half of the reminder cues (i.e., sounds followed by the first syllable of the associated words, similar to the cued recall procedure) were presented in SWS and the other half in S2. SWS-cues and S2-cues were presented blockwise, i.e., either all SWS-cues first and then all S2-cues or vice versa. The sleep stage order was counterbalanced across subjects. Each sound (45 dB) was presented only once for 2.9 s and was then accompanied by the first syllable of the word for 1.5 s. After a 7 s interval, the next reminder cue was presented. Reactivation could be paused whenever signs of arousal or changes in sleep stage were detected, and restarted upon detection of the corresponding sleep stage.

### Control task

At the end of the experiment, participants were asked to report if they had heard any sounds or words while they were sleeping (‘Heard/not-heard task’). The 30 sounds plus first syllables were presented again and subjects had to indicate if they think that they had received those stimuli during sleep or not.

### Sleep recording

Standard polysomnography was obtained including electroencephalographic (EEG), electromyographic (EMG), and electrooculographic (EOG) recordings with BrainAmp amplifiers (Brain Products, Munich, Germany). EEG was recorded from six scalp electrodes (F3, F4, C3, C4, P3 and P4 according to the International 10–20 System) and two electrodes on the left and right mastoids were used as combined reference. Sampling rate was 200 Hz and data was bandpass-filtered between 0.16 and 35 Hz. Polysomnographic recordings were scored offline as wake, stage 1, stage 2, stages 3 and 4 (SWS), and REM sleep according to standard criteria by Rechtschaffen & Kales^18^.

### Sleep EEG analyses

EEG data was analyzed using custom-made codes in MATLAB 2020a (Mathworks). Data was zero-phase filtered in the bands from 1 to 35 Hz (Butterfilter). Each reactivation event was cut into 12 s trials (–7 to 5 s with t = 0 referring to the syllable cue onset). For one subject, reactivation markers were missing, thus leaving 19 subjects for the EEG analysis. From the remaining subjects, in total 15 trials with artifacts were excluded from the analysis. Trials were averaged per subject and finally a grand average for each condition was calculated (SWS-cueing vs. S2-cueing).

To evaluate the braińs response to auditory cueing, evoked response potentials (ERP) were calculated aligned to the syllable cue onset and averaged for each condition. ERP amplitude was calculated from peak-to-peak and averaged for each subject.

Time-frequency analyses were performed for each sleep stage separately. Power was calculated relative to a baseline from –4 and –3 s (i.e., right before the sound onset). This processing resulted in the relative power of each frequency at each time point. ERP and time-frequency analyses were performed on central electrodes (i.e., average of C3 and C4).

We further detected discrete slow oscillations (SO) and sleep spindles within three periods of interest: 1) during the cueing period (“React”), 2) a period immediately before the cueing period (“Pre”) and 3) a period immediately after the cueing period (“Post”). For those subjects, who had the entire cueing session without arousals and interruptions, the session took 2.85 min, i.e., ∼3 min (for the 15 cues). Thus, we took 3 min as a standard and considered 3 min before the reactivation session and 3 min after the reactivation as pre-cueing and post-cueing periods, respectively. Since some subjects had interruptions in the cueing period, we only considered spindles and SO in those epochs with cueing markers for the cueing period.

The detection algorithm we used to identify SO and spindles is based on the methods of Mölle and collaborators^53^. A SO was detected when the peak-to-peak amplitude was larger than 75 µV in the filtered EEG (0.05-3.5 Hz). For fast spindle detection, the EEG was filtered between 12-15 Hz, then we calculated the root mean square (RMS) and applied a moving average of 200 ms. Using this smoothed RMS, a spindle was detected when this value was above the threshold of 1.5 standard deviation values from the mean, between 0.5-3 s. We also detected slow spindles, however, since there were no differences in any of the measures, we do not further report these findings and all reported analyses refer to fast spindles.

To examine the coupling of fast spindles to SO, we identified SO-spindle complexes for the three periods of interest defined above. We calculated the density of SO-spindle complexes (events/minute) during “Pre”, “React” and “Post” periods. SOs, fast spindles and SO-Spindle complexes density was calculated for the average of central, frontal and parietal channels. In the main text, we only report results of central channels for SO and SO-Spindle complexes, and parietal channels for spindles. However, analyses for the other channels show very similar patterns of findings (see Supplementary Information).

### Statistics

Statistical analyses for behavioral and EEG analyses were performed in SPSS 28.0.1.1 and Matlab 2020a (Mathworks). Memory change (i.e., number of correct responses at testing minus the number of correct responses at training) was used as measure for memory performance. Statistical comparisons between SWS-cueing and S2-cueing conditions were done with paired-sample t-tests for memory change, learning measures, the interference task, and the different sleep parameters such as ERP amplitude. Additionally, a repeated-measures ANOVA with within-subjects factor “sleep stage” and between-subjects factor “order” was performed to check for the effect of the reactivation order (first reactivation in S2 vs. first reactivation in SWS).

For the EEG analyses, repeated-measures ANOVAs were applied to compare SO density, fast spindles and SO-spindle complexes between SWS and S2 across the different periods of interest (“Pre”, “React” and “Post”). Bonferroni correction for multiple comparisons was performed.

A value of P < 0.05 was considered significant. Effect size analyses were computed in Gpower 3.1. Bayesian analyses were performed in JASP 0.17.3.0 to formally test whether, in the absence of significant differences between conditions, the data support the null hypothesis.

## Acknowledgments

This work was funded by a joint collaboration grant from the Deutsche Forschungsgemeinschaft (DFG) to SD (DI 1866-2-1) and CONICET/MINCyT to CF (Resolución D. No 4427), as well as a doctoral research fellowship from the Deutscher Akademischer Austauschdienst (DAAD) to JC. We would like to thank Karsten Rauss for support with the Bayesian statistics. We would further like to mention that different findings from the present study had previously been reported in conference presentations (i.e., stronger cueing effects for SWS than S2). These findings were erroneous due to switched assignment of conditions in some subjects. This error has now been corrected and the present findings have been validated in additional subjects.

## Authors contributions

SD, CF, and JC conceived the experiments, JC collected the data, JC, CB, and SD analyzed the data, all authors wrote and reviewed the manuscript.

## Data availability

The dataset supporting the findings of this study are available from the corresponding author upon request.

**Correspondence** and requests for materials should be addressed to Susanne Diekelmann

## References

1. Diekelmann, S. & Born, J. The memory function of sleep. Nat. Rev. Neurosci. 11, 114–126 (2010).

2. Rasch, B. & Born, J. About sleep’s role in memory. Physiol. Rev. 93, 681–766 (2013).

3. Stickgold, R. Neuroscience: A memory boost while you sleep. Nature 444, 559–560 (2006).

4. Paller, K. A., Creery, J. D. & Schechtman, E. Memory and Sleep: How Sleep Cognition Can Change the Waking Mind for the Better. Annu. Rev. Psychol. 72, 123–130 (2021).

5. Ackermann, S. & Rasch, B. Differential effects of non-REM and REM sleep on memory consolidation? Curr. Neurol. Neurosci. Rep. 14, (2014).

6. Gais, S. & Born, J. Declarative memory consolidation: Mechanisms acting during human sleep. Learn. Mem. 11, 679–685 (2004).

7. Mölle, M., Eschenko, O., Gais, S., Sara, S. J. & Born, J. The influence of learning on sleep slow oscillations and associated spindles and ripples in humans and rats. Eur. J. Neurosci. 29, 1071–1081 (2009).

8. Ngo, H. V., Fell, J. & Staresina, B. Sleep spindles mediate hippocampal-neocortical coupling during long-duration ripples. Elife 9, 1–18 (2020).

9. Staresina, B. P. et al. Hierarchical nesting of slow oscillations, spindles and ripples in the human hippocampus during sleep. Nat. Neurosci. 18, 1679–1686 (2015).

10. Helfrich, R. F., Mander, B. A., Jagust, W. J., Knight, R. T. & Walker, M. P. Old Brains Come Uncoupled in Sleep: Slow Wave-Spindle Synchrony, Brain Atrophy, and Forgetting. Neuron 97, 221–230 (2018).

11. Muehlroth, B. E. et al. Precise Slow Oscillation–Spindle Coupling Promotes Memory Consolidation in Younger and Older Adults. Sci. Rep. 9, 1–15 (2019).

12. Ladenbauer, J. et al. Promoting sleep oscillations and their functional coupling by transcranial stimulation enhances memory consolidation in mild cognitive impairment. J. Neurosci. 37, 7111–7124 (2017).

13. Niknazar, M., Krishnan, G. P., Bazhenov, M. & Mednick, S. C. Coupling of thalamocortical sleep oscillations are important for memory consolidation in humans. PLoS One 10, 1–14 (2015).

14. Demanuele, C. et al. Coordination of slow waves with sleep spindles predicts sleep-dependent memory consolidation in schizophrenia. Sleep 40, 1–10 (2017).

15. Lewis, P. A. & Durrant, S. J. Overlapping memory replay during sleep builds cognitive schemata. Trends Cogn. Sci. 15, 343–351 (2011).

16. Genzel, L., Kroes, M. C. W., Dresler, M. & Battaglia, F. P. Light sleep versus slow wave sleep in memory consolidation: A question of global versus local processes? Trends Neurosci. 37, 10–19 (2014).

17. Cox, R., Mylonas, D. S., Manoach, D. S. & Stickgold, R. Large-scale structure and individual fingerprints of locally coupled sleep oscillations. Sleep 41, 1–15 (2018).

18. Rechtschaffen A, Kales, A. *A Manual of Standardized Terminology, Technique and Scoring System for Sleep Stages of Human Sleep*. (Brain Information Service, Brain Information Institute, 1968).

19. Wilhelm, I. et al. Sleep selectively enhances memory expected to be of future relevance. J. Neurosci. 31, 1563–1569 (2011).

20. Marshall, L., Helgadóttir, H., Mölle, M. & Born, J. Boosting slow oscillations during sleep potentiates memory. Nature 444, 610–613 (2006).

21. Huber, R., Ghilardi, M. F., Massimini, M. & Tononi, G. Local sleep and learning. Nature 430, 78–81 (2004).

22. Mölle, M., Marshall, L., Gais, S. & Born, J. Learning increases human electroencephalographic coherence during subsequent slow sleep oscillations. Proc. Natl. Acad. Sci. U. S. A. 101, 13963–13968 (2004).

23. Gais, S., Mölle, M., Helms, K. & Born, J. Learning-dependent increases in sleep spindle density. J. Neurosci. 22, 6830–4 (2002).

24. Ruch, S. et al. Sleep stage II contributes to the consolidation of declarative memories. Neuropsychologia 50, 2389–2396 (2012).

25. Schabus, M. et al. Sleep spindles and their significance for declarative memory consolidation. Sleep 27, 1479–1485 (2004).

26. Seeck-Hirschner, M. et al. Declarative Memory Performance Is Associated With the Number of Sleep Spindles in Elderly Women. Am. J. Geriatr. Psychiatry 20, 782–788 (2012).

27. Hahn, M. A., Heib, D., Schabus, M., Hoedlmoser, K. & Helfrich, R. F. Slow oscillation-spindle coupling predicts enhanced memory formation from childhood to adolescence. Elife 9, 1–21 (2020).

28. Hu, X., Cheng, L. Y., Chiu, M. H. & Paller, K. A. Promoting memory consolidation during sleep: A meta-analysis of targeted memory reactivation. Psychol. Bull. 146, (2020).

29. Klinzing, J. G. & Diekelmann, S. Chapter 31 – Cued Memory Reactivation: A Tool to Manipulate Memory Consolidation During Sleep. in Handbook of Sleep Research (ed. Dringenberg, H. C. B. T.-H. of B. N.) **30**, 471–488 (Elsevier, 2020).

30. Oudiette, D. & Paller, K. A. Upgrading the sleeping brain with targeted memory reactivation. Trends Cogn. Sci. 17, 142–149 (2013).

31. Diekelmann, S., Büchel, C., Born, J. & Rasch, B. Labile or stable: Opposing consequences for memory when reactivated during waking and sleep. Nat. Neurosci. 14, 381–386 (2011).

32. Schreiner, T. & Rasch, B. Boosting vocabulary learning by verbal cueing during sleep. Cereb. Cortex 25, 4169–4179 (2015).

33. Farthouat, J., Gilson, M. & Peigneux, P. New evidence for the necessity of a silent plastic period during sleep for a memory benefit of targeted memory reactivation. Sleep Spindl. Cortical Up States 1, 14–26 (2017).

34. Cairney, S. A., Durrant, S. J., Hulleman, J. & Lewis, P. A. Targeted memory reactivation during slow wave sleep facilitates emotional memory consolidation. Sleep 37, 701–707 (2014).

35. Diekelmann, S., Biggel, S., Rasch, B. & Born, J. Offline consolidation of memory varies with time in slow wave sleep and can be accelerated by cuing memory reactivations. Neurobiol. Learn. Mem. 98, 103–111 (2012).

36. Diekelmann, S., Born, J. & Rasch, B. Increasing explicit sequence knowledge by odor cueing during sleep in men but not women. Front. Behav. Neurosci. 10, 1–11 (2016).

37. Forcato, C. et al. Reactivation during sleep with incomplete reminder cues rather than complete ones stabilizes long-term memory in humans. *Commun*. Biol. 3, 1–13 (2020).

38. Rasch, B., Büchel, C., Gais, S. & Born, J. Odor Cues During Slow-Wave Sleep Prompt Declarative Memory Consolidation. Science (80-.). 315, 1426–1429 (2007).

39. Cairney, S. A., Guttesen, A. á. V., El Marj, N. & Staresina, B. P. Memory Consolidation Is Linked to Spindle-Mediated Information Processing during Sleep. Curr. Biol. 28, 948–954 (2018).

40. Rudoy, J. D., Voss, J. L., Westerberg, C. E. & Paller, K. A. Strengthening individual memories by reactivating them during sleep. Science (80-.). 326, 1079 (2009).

41. Schönauer, M., Geisler, T. & Gais, S. Strengthening Procedural Memories by Reactivation in Sleep. J. Cogn. Neurosci. 26, 143–153 (2014).

42. Schreiner, T., Doeller, C. F., Jensen, O., Rasch, B. & Staudigl, T. Theta Phase-Coordinated Memory Reactivation Reoccurs in a Slow-Oscillatory Rhythm during NREM Sleep. Cell Rep. 25, 296–301 (2018).

43. Ritter, S. M., Strick, M., Bos, M. W., Van Baaren, R. B. & Dijksterhuis, A. Good morning creativity: Task reactivation during sleep enhances beneficial effect of sleep on creative performance. J. Sleep Res. 21, 643–647 (2012).

44. Neumann, F., Oberhauser, V. & Kornmeier, J. How odor cues help to optimize learning during sleep in a real life-setting. Sci. Rep. 10, 1–8 (2020).

45. Vidal, V. et al. Odor cueing during sleep improves consolidation of a history lesson in a school setting. Sci. Rep. 12, 1–7 (2022).

46. Göldi, M. & Rasch, B. Effects of targeted memory reactivation during sleep at home depend on sleep disturbances and habituation. *npj Sci*. Learn. 4, 1–7 (2019).

47. Wick, A. & Rasch, B. Targeted memory reactivation during slow-wave sleep vs. sleep stage N2: no significant differences in a vocabulary task. Learn. Mem. 30, 192–200 (2023).

48. Carbone, J. et al. The effect of zolpidem on targeted memory reactivation during sleep. Learn. Mem. 28, 307–318 (2021).

49. Ngo, H. V. V., Martinetz, T., Born, J. & Mölle, M. Auditory closed-loop stimulation of the sleep slow oscillation enhances memory. Neuron 78, 545–553 (2013).

50. Groch, S., Schreiner, T., Rasch, B., Huber, R. & Wilhelm, I. Prior knowledge is essential for the beneficial effect of targeted memory reactivation during sleep. Sci. Rep. 7, 1–7 (2017).

51. Schreiner, T., Göldi, M. & Rasch, B. Cueing vocabulary during sleep increases theta activity during later recognition testing. Psychophysiology 52, 1538–1543 (2015).

52. Mikutta, C. et al. Phase-amplitude coupling of sleep slow oscillatory and spindle activity correlates with overnight memory consolidation. J. Sleep Res. 28, 1–6 (2019).

53. Mölle, M., Bergmann, T. O., Marshall, L. & Born, J. Fast and slow spindles during the sleep slow oscillation: Disparate coalescence and engagement in memory processing. Sleep 34, 1411–1421 (2011).

