## Supplementary information for "Targeted memory reactivation is not more effective during slow wave sleep than sleep stage 2"

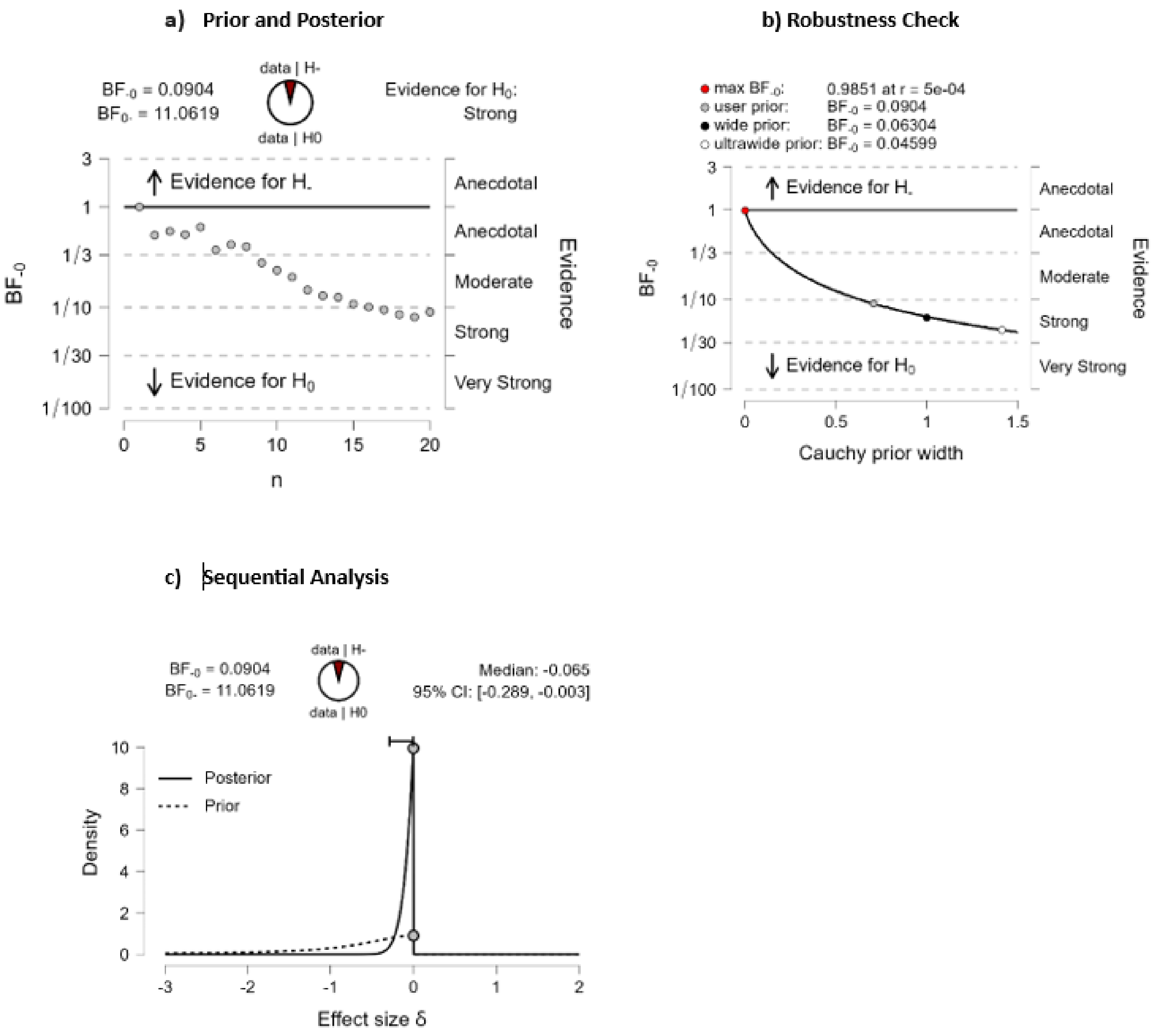


**Supplementary Figure S1. Bayesian statistics on memory performance. a.** Prior and posterior distributions and Bayes Factors for the unidirectional test of the hypothesis that stimulation during SWS should yield higher memory performance than during S2. The resulting Bayes Factors indicate that the null model is 11 times more likely when assuming JASP's default prior (Cauchy distribution with scale 0.707). **b.** A robustness check across different prior widths indicates that a more narrow prior (indicating an a priori assumption of finding relatively small effects) would still result in moderate support of the null model. Under no conditions do the data offer support for the alternative hypothesis. **c.** A sequential analysis across subjects indicates asymptotic development of the Bayes Factor, further supporting the notion that there is no difference between stimulation during S2 and SWS.

**Supplementary Table S1. Sleep oscillatory activity for central, frontal and parietal channels**

| **C Channel** | **Pre S2** | **React S2** | **Post S2** | **Pre SWS** | **React SWS** | **Post SWS** |
|---|---|---|---|---|---|---|
| SO density | 2.86 ± 0.85 | 3.97 ± 0.91 | 4.63 ±0.73 | 16.68 ±1.75 | 22.58 ±2.11 | 24.93 ± 1.82 |
| Spindle density | 1.51 ± 0.45 | 1.67 ± 0.26 | 2.05 ± 0.40 | 3.25 ± 0.53 | 2.59 ± 0.37 | 2.44 ± 0.37 |
| SO-Spindle complexes density | 0.26 ± 0.09 | 0.36 ± 0.07 | 0.75 ± 0.15 | 1.28 ± 0.15 | 0.89 ± 0.14 | 1.14 ± 0.19 |
| **F channel** | | | | | | |
| SO density | 4.96 ± 1.51 | 6.15 ± 1.51 | 7.91 ± 1.36 | 25.09 ± 2.60 | 32.09 ± 2.78 | 32.70 ± 2.44 |
| Spindle density | 1.32 ± 0.46 | 1.39 ± 0.28 | 2.11 ± 0.47 | 3.42 ± 0.49 | 3.33 ± 0.48 | 3.32 ± 0.54 |
| SO-Spindle complexes density | 0.21 ± 0.07 | 0.42 ± 0.16 | 0.91 ± 0.21 | 1.68 ± 0.22 | 1.24 ± 0.24 | 1.35 ± 0.24 |
| **P Channel** | | | | | | |
| SO density | 2.33 ± 0.92 | 3.04 ± 0.95 | 3.09 ± 0.54 | 13.86 ± 1.60 | 19.41 ± 1.96 | 21.79 ± 1.83 |
| Spindle density | 1.14 ± 0.42 | 1.53 ± 0.23 | 2.54 ± 0.50 | 3.61 ± 0.53 | 2.96 ± 0.45 | 2.86 ± 0.46 |
| SO-Spindle complexes density | 0.19 ± 0.07 | 0.29 ± 0.09 | 0.42 ± 0.10 | 1.16 ± 0.15 | 0.73 ± 0.13 | 0.98 ± 0.14 |

Slow oscillation (SO) density, spindle density and SO-spindle complexes density during three periods of interest: cue presentation (“React”, 3 min), pre-cueing (“Pre”, 3 min), and post-cueing (“Post”, 3 min). Means ± SEM are shown for SWS and S2, separately for central (C) frontal (F) and parietal (P) channels.

**Statistical analysis of sleep oscillatory activity**

We performed repeated-measures ANOVAs with the within-subjects factors “Sleepstage” (S2 and SWS), and “Period” (Pre, React and Post). Uncorrected p values are reported in the following. When Bonferroni correction for multiple comparisons is applied, p values <0.005 remain significant.

For the central channels, the repeated-measures ANOVA for SO density revealed significant main effects for Sleepstage ($p<0.001$) and Period ($p<0.001$), as well as for the interaction Sleepstage*period ($p<0.001$). Post-hoc Bonferroni Tests yielded significant differences between Pre S2 and Pre SWS ($p<0.001$), between React S2 and React SWS ($p<0.001$), and between Post S2 and Post SWS ($p<0.001$). Moreover, there were significant differences when comparing SWS periods, particularly for Pre SWS and React SWS ($p<0.001$), and Pre SWS and Post SWS ($p<0.001$).

Repeated-measures ANOVA for spindle density revealed no significant main effect for Sleepstage ($p=0.058$) or Period (0.654), but a significant interaction effect Sleepstage*period ($p=0.049$). No significant Post-hoc effects were found.

Repeated-measures ANOVA for SO-spindle complexes revealed significant main effects for Sleepstage ($p<0.001$) and Period ($p<0.001$), as well as for the interaction Sleep Stage*period ($p<0.001$). Post-hoc Bonferroni Tests yielded significant differences between Pre S2 and Pre SWS ($p<0.001$), between React S2 and React SWS ($p<0.001$), and between Post S2 and Post SWS ($p<0.001$). Moreover, there were significant differences when comparing between SWS periods, particularly for Pre SWS and React SWS ($p<0.001$), and Pre SWS and Post SWS ($p<0.001$).

For the frontal channels, the repeated-measures ANOVA for SO density revealed significant main effects for Sleepstage ($p<0.001$) and Period ($p<0.001$), as well as for the interaction Sleepstage*period ($p=0.005$). Post-hoc Bonferroni Tests yielded significant differences between Pre S2 and Pre SWS ($p<0.001$), between React S2 and React SWS ($p<0.001$), and between Post S2 and Post SWS ($p<0.001$). Moreover, there were significant differences when comparing SWS periods, particularly for Pre SWS and React SWS ($p<0.001$), and Pre SWS and Post SWS ($p<0.001$).

Repeated-measures ANOVA for spindle density revealed no significant main effect for Period ($p=0.338$) but for Sleepstage ($p=0.003$) and no significant interaction effect Sleepstage*period ($p=0.338$). Post-hoc Bonferroni Tests yielded significant differences between Pre S2 and Pre SWS ($p=0.033$), and between React S2 and React SWS ($p=0.043$).

Repeated-measures ANOVA for SO-spindle coupling revealed significant main effects for Sleepstage ($p<0.001$) and Period ($p=0.034$), as well as for the interaction Sleepstage*period ($p=0.004$). Post-hoc Bonferroni Tests yielded significant differences between Pre S2 and Pre SWS ($p<0.001$), and between React S2 and React SWS ($p=0.022$). Moreover, there were significant differences when comparing S2 periods, particularly for Pre S2 and Post S2 ($p=0.003$)

For parietal channels, repeated-measures ANOVA for SO density revealed significant main effects for Sleepstage ($p<0.001$) and Period ($p<0.001$), as well as for the interaction Sleepstage*period ($p<0.001$). Post-hoc Bonferroni Tests yielded significant differences between Pre S2 and Pre SWS ($p<0.001$), between React S2 and React SWS ($p<0.001$), and between Post S2 and Post SWS ($p<0.001$). Moreover, there were significant differences when comparing SWS periods, particularly for Pre SWS and React SWS ($p<0.001$), and Pre SWS and Post SWS ($p<0.001$).

Repeated-measures ANOVA for Spindle density revealed no significant main effect for Period ($p=0.153$) but for Sleep stage ($p=0.019$) and a significant interaction effect Sleepstage*period ($p=0.002$). Post-hoc Bonferroni Tests yielded significant differences between Pre S2 and Pre SWS ($p=0.007$). Moreover, there were significant differences when comparing S2 periods, particularly between Pre S2 and Post S2 ($p=0.004$).

Repeated-measures ANOVA for SO-spindle coupling revealed no significant main effect for Period ($p=0.061$), but for Sleepstage ($p<0.001$), as well as for the interaction Sleepstage*period ($p=0.021$). Post-hoc Bonferroni Tests yielded significant differences between Pre S2 and Pre SWS ($p<0.001$), and between Post S2 and Post SWS ($p=0.016$). Moreover, there were significant differences when comparing SWS periods, particularly for Pre SWS and react SWS ($p=0.012$).
